# Comparing Fecal, Saliva and Chicha Microbiomes Between Mothers and Children in an Indigenous Ecuadorian Cohort

**DOI:** 10.1101/2020.10.02.323097

**Authors:** Eric Adams, Andrew Oliver, Alexandria Gille, Nadia Alaniz, Carolina Jaime, John Patton, Katrine Whiteson

## Abstract

Recent research has elucidated many factors which play a role in the development and composition of human microbiomes. In this study we briefly examine the microbiomes of saliva and fecal samples from 71 indigenous individuals, and chicha samples from 28 single family households in a remote community in the Ecuadorian Amazon. Fecal and saliva samples were collected at two separate time points whereas chicha samples were collected at four time points, once each day of the fermentation process. In total 324 samples were collected: 113 saliva, 108 chicha, and 103 fecal. Microbial composition and diversity were assessed using shotgun metagenome sequence data. Chicha samples were found to be nearly entirely composed of the order *Lactobacillales*, accounting for 90.1% of the relative abundance. Saliva samples also contained a high relative abundance of *Lactobacillales* (31.9%) as well as being composed of *Neisseriales* (12.8%), *Actinomycineae* (8.7%), *Bacteroidales* (7.0%), *Clostridiales* (6.8%), *Micrococcineae* (6.5%), and *Pasteurellales* (6.0%). Fecal samples were largely composed of the three orders *Clostridiales* (33.7%), *Bacteroidales* (21.9%), and *Bifidobacteriales* (16.5%). Comparison of α-diversity, as calculated by Shannon’s Diversity Index, in mothers and their offspring showed no significant difference between the two groups in either fecal or saliva samples. Comparison of β-diversity in fecal and saliva samples, as calculated by the Bray-Curtis Dissimilarity measure, within household units and between differing households showed that members of the same household were significantly less dissimilar to each other than to members of other households in the community. Average microbiome composition for individuals within fecal and saliva samples was assessed to determine the impact of an individual’s household on the composition of their microbiome. Household was determined to have a significant impact on both fecal and oral microbiome compositions.

## INTRODUCTION

In recent decades there has been a growing abundance of research into the microbial organisms which live in and around people and animals. These malleable communities of microorganisms are referred to as microbiomes and they interact with host organisms in a number of ways, many of the details and significance of which are still being discovered. Studies of human microbiomes have shown that they are involved in a number of physiological processes such as nutrient metabolism, immune system development, and resistance against invading pathogens.^1,2^ A number of researchers are investigating the factors which influence the development of, and differences between, various microbiomes. Some work is more focused on factors which influence early development or the relative contributions of genetics and environment, while others examine the impacts of lifestyle, economic, and dietary differences on microbiome composition.^1–5^ Some efforts have even been made to assemble models of community structure and microbial transfer networks from microbiome data in isolated communities.^6,7^

The advances in this field of research are due to the ever-increasing affordability of genomic sequencing and the increasing power of computational analysis tools allowing for rapid metagenomic analysis of microbial communities. Metagenomics is a powerful method of investigating the composition of microbiomes because it involves the collection and amplification of all the genomic DNA present in a collected sample. This presents a less biased view of the microbes present because it is not limited to the organisms which can be cultured in a lab. Many metagenomic studies into microbiomes rely on amplification of variable regions of the bacterial gene for the 16S small-subunit ribosomal RNA which allows for taxonomic and phylogenetic classifications of samples. This study utilized shotgun metagenomic sequencing, where instead of amplifying a single region of the genome, as in 16S rRNA sequencing, shotgun metagenomics instead shears the DNA in a sample into fragments which are independently amplified and sequenced. This enables both taxonomic classification as well as information on functional genes within the microbiome, providing a picture of which species are present and the metabolic processes they encode.^8^ The widespread use of both of these sequencing methods for microbial analysis has led to a number of databases containing the sequence data of human microbiomes. A common feature of many recent studies into the microbiome involves the comparison of isolated or semi-isolated indigenous groups to more industrialized western cohorts.^9,10^

The subjects of this research are a group of indigenous people living in Conambo, a village along the Conambo river, deep within the jungles of Ecuador (**Fig. 1A**). This group has been extensively studied and characterized for many years by anthropologists led by Dr. John Patton and Brenda Bowser, and is a very isolated community with little outside contact.^11–13^ The small village of just under 200 people is centered around a community building, school, and small dirt airstrip that the people maintain. As there are no roads to the village and access by boat is impossible upriver while closed at the border with Peru downriver, a small single propeller plane is the only way that this village is reached by the missionaries and anthropologists with which the villagers occasionally interact. The community center located along the airstrip is where the people of the village hold communal meetings, ceremonies, and celebrations. There is also a radio which is their only means to contact the outside world. An event called a “ minga” is held when a member of the community needs help on a task such as clearing a garden, collecting thatching for a roof, or some other labor-intensive work; but there is otherwise little contact between some of the more spread apart households. The individual households are relatively far from one another, often requiring treks of a few kilometers through hills and dense jungle. Households in the village are composed of a male head of household, his wife, or wives, as well as any children they have. Villagers in this area practice what is referred to as self-sufficient horticultural foraging. As there is no way for them to purchase or trade for food and crops, they subsist by hunting and foraging as well as crop gardens maintained by the women of each individual household.

**Figure 1.**
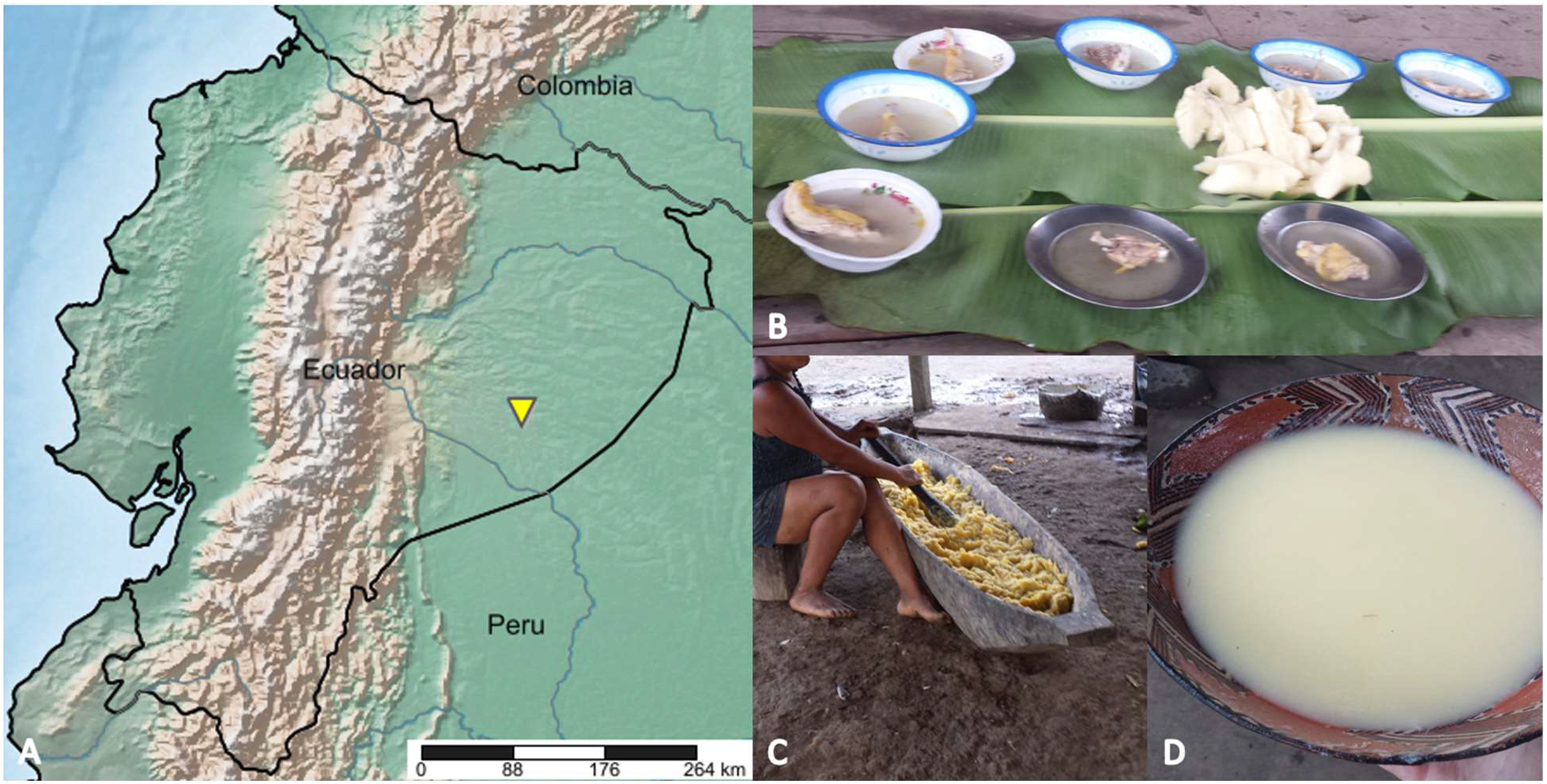
(A) Location of Conambo (yellow triangle) within the borders of Ecuador; map generated using simplemappr.net (B) A meal set out for visiting guests consisting of stewed meat and boiled plantains that is representative of the normal dietary intake (C) Traditional preparation method for the yuca mash which is fermented to produce chicha (D) A serving of chicha in a traditional clay bowl made in the village

The normal diet of the people includes a stew of wild game, (mostly monkey, tapir; peccary and large rodents), fish and various birds, served alongside boiled yuca or plantains (**Fig. 1B**). A beverage made from fermented yuca root, known as “ chicha,” also serves as a major source of calories for these people. The chicha is made by the maternal head of the household, sometimes with the assistance of her daughters by repeated mastication of the boiled yuca root in a large wooden trough (**Fig. 1C**). The production process of the mash is likely what inoculates it with fermentative bacteria from saliva.^14^ After the mash is prepared, it is covered in banana leaves and allowed to ferment for four days before it is mixed with water and served in a traditional ceramic bowl (**Fig. 1D**). Interestingly, in this village the people almost exclusively drink chicha as opposed to water or other liquids, indicating that there may be a high degree of transfer between oral and chicha microbiomes.

This study’s purpose is to compare the diversity levels of gut microbiota in mothers and their offspring to determine if, as shown in prior studies, they show highly similar levels of diversity.^2^ The dissimilarity of maternal microbiomes to those of their offspring will also be compared to the dissimilarity of the community at large. These two factors are intended to show if there is a high degree of similarity between the microbiomes of mother-offspring groups and whether they are distinct from the people around them, as there is an abundance of research showing the maternal influence on the development of the microbiome.^2,3^ Given the highly similar environmental conditions of this indigenous group, a more detailed analysis of the functional data could determine the degree of maternal and family influence on microbiome composition.

## MATERIALS AND METHODS

### Sample collection and genomic DNA extraction

The sets of saliva, chicha, and fecal samples were collected in the field by anthropology graduate students working under Dr. John Patton from CSU Fullerton as a part of a longer, ongoing ethnographic study. Saliva samples were collected at two intervals, coinciding with collection of the first and last chicha samples. Chicha samples were collected at four intervals across the four days of the fermentation process. Fecal samples were collected at two intervals with collection being performed by the subject after verbal explanation of sampling method. A total of 113 saliva, 108 chicha, and 103 fecal samples were collected in June 2018 and stored using DNA/RNA Shield swab collection tubes (Zymo Research, Cat# R1107). The samples were gathered from a total of 71 individuals in 28 households, encompassing every adult female living in the community as well as their offspring up to age seven residing with them. Samples were not collected from adult males in the community. The sample collection tubes were transported to CSU Fullerton and kept in a freezer at -20°C until transfer to UC Irvine for extraction and processing.

Microbial genomic DNA was extracted from each of the samples using the ZymoBIOMICS 96 DNA Kit (Zymo Research, Cat# D4309) according to the manufacturer’s instructions. This included 5 × 1-minute bed-beating steps using a FastPrep-24 (MP Biomedicals, Cat# SKU 116004500) at maximum speed. The quantity of DNA in each sample was fluorescently measured with the Quant-iT PicoGreen dsDNA Assay Kit (ThermoFicher, Cat# P11496) using a Synergy H1 Microplate reader (BioTek, Cat # BTH1M).

### Library preparation and sequencing

Sequence libraries were prepared from the extracted DNA samples using the Nextera DNA Flex Library Prep Kit (Illumina, Cat. # 20018705) following a low volume variation of the standard protocol. Normalization of DNA concentration prior to creation of sequence libraries is not required with the Nextera Flex kit which was one of the reasons that this method was selected. Samples were prepared for PCR with Kapa HiFi HotStart ReadyMix (Roche, Cat # 07958935001) using custom primers ordered from Integrated DNA Technologies. All of the PCR steps were performed in an Eppendorf Mastercycler Nexus Gradient (Eppendorf, Cat # 2231000665) using the standard thermal cycles as described in the Nextera Flex protocol. The resulting sequence fragments were analyzed on an Agilent Bioanalyzer to determine fragment length distribution (Agilent, Cat # G2939BA). Sequence libraries were pooled based on DNA concentration as determined by the Bioanalyzer. After pooling, the libraries were sent to Novogene Co., Ltd. for sequencing on an Illumina HiSeq4000.

### Sequence assembly and statistical analysis

Sequence reads were first quality filtered to a Q_phred_ score of greater than 30. Quality filtered sequences were then mapped to references of the human genome, mitochondria, and chloroplasts; matching sequences were removed from further analysis. The remaining sequences were mapped against marker genes as part of the MiDAS pipeline to generate a taxonomic abundance table.^15^ Samples were rarified to even levels of 300 MiDAS marker gene hits per sample for all statistical analysis. Statistical analysis was completed in R with α-diversity calculated using the Shannon Diversity Index and β-diversity calculated using the Bray-Curtis dissimilarity measure using the vegan package. PERMANOVA was performed using the vegan package and allowing for 999 permutations, the Kruskal-Wallis one-way analysis of variance test was used to determine the significance of diversity and dissimilarity indexes, and plots were generated in R using ggplot2.

## RESULTS

A total of 324 samples from 71 individuals were collected in the field and stored in DNA/RNA shield solutions. The samples, consisting of either saliva, feces, or chicha, were extracted for genomic DNA which was then fragmented and prepared into libraries for shotgun sequencing on an Illumina HiSeq platform. Extractions yielded an average DNA concentration of 44.02 ng/μL from saliva samples, 116.47 ng/μL from fecal samples, and 6.11 ng/μL from chicha samples. Preparation of the DNA sequencing libraries yielded a fragment distribution with an average fragment length of 489 bp. Sequencing of the prepared libraries yielded 505,329,634 raw reads. These raw sequences were quality screened by Novogene Co., Ltd. to remove reads that contained adapters, had N > 10%, or a Q_score_ ≤ 5, resulting in 505,140,753 initially filtered reads.

The initially filtered sequence reads from Novogene Co., Ltd. were further quality filtered to a Phred score of 30 or greater using Prinseq, though most sequence reads had Q-scores closer to 40. Quality filtered sequence reads were mapped against marker genes using the MiDAS pipeline, yielding a taxonomic abundance table for each sample. A majority of sequence reads could not be identified as marker genes using MiDAS and approximately 75,000 sequences were classified into taxonomic abundance tables from the ∼250 million filtered sequence reads.

The taxonomic classifications generated from MiDAS were sorted according to the sample type and the relative abundance of the bacterial orders were plotted for chicha, fecal, and saliva samples (**Fig. 2**). Chicha samples consisted mainly of the order *Lactobacillales* which accounted for 90.1% of the relative abundance, with the remainder consisting of *Rhodospirillales* (4.3%) and a number of orders with low abundances. Fecal samples were largely composed of three major orders: *Clostridiales*, accounting for 33.7% of the relative abundance; *Bacteroidales*, accounting for 21.9% of the relative abundance; and *Bifidobacteriales*, accounting for 16.5% of the relative abundance. Saliva samples showed a more equal distribution of relative abundances across the most abundant orders. The most abundant order in the saliva samples was the *Lactobacillales*, which accounted for 31.9% of the relative abundance. The next most abundant order in the saliva samples was *Neisseriales*, comprising 12.8% of the relative abundance. There were also relatively equal abundances of the orders *Actinomycineae* (8.7%), *Bacteroidales* (7.0%), *Clostridiales* (6.8%), *Micrococcineae* (6.5%), and *Pasteurellales* (6.0%) present in the saliva samples.

**Figure 2.**
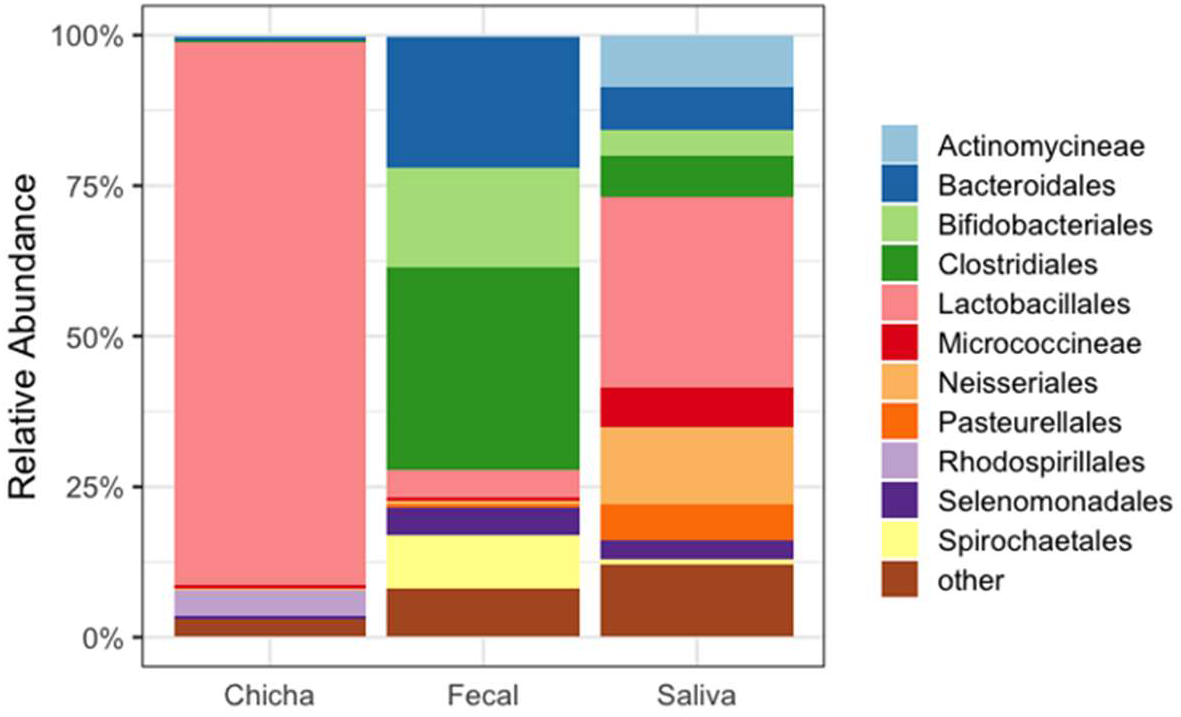
Relative abundances of microbial orders observed in each of the three sample types. The chicha samples were almost entirely dominated by the *Lactobacillales* order. The saliva samples also showed a high abundance of the *Lactobacillales*, as well as *Neisseriales* and *Actinomycineae*. Fecal samples are mainly dominated by the orders *Clostridiales, Bacteroidales*, and *Bifidobacteriales*. Number of individuals and samples represented in each sample type: Chicha (31 individuals, 100 samples), Fecal (49 individuals, 78 samples), Saliva (42 individuals, 62 samples).

The level of α-diversity in mother and child samples was determined using the Shannon Diversity Index, which provides a characterization of both the abundance and richness of the species present in a sample (**Fig. 3**). The Shannon Diversity for the microbiomes in fecal samples had a median value of 2.70 for mothers and 2.73 for children. Saliva samples from both groups were found to have a higher Shannon’s Diversity Index value at 3.64 for mothers and 3.71 for children. The Kruskal-Wallis one-way analysis of variance was used to determine if there was a significant difference between the diversity measures of mothers and their offspring. The diversity measures in both fecal and saliva samples between mothers and children did not significantly differ from one another (p = 0.33 and p = 0.58 respectively). Further information can be gained by not only looking at similarity in diversity, but also dissimilarity in species presence.

**Figure 3.**
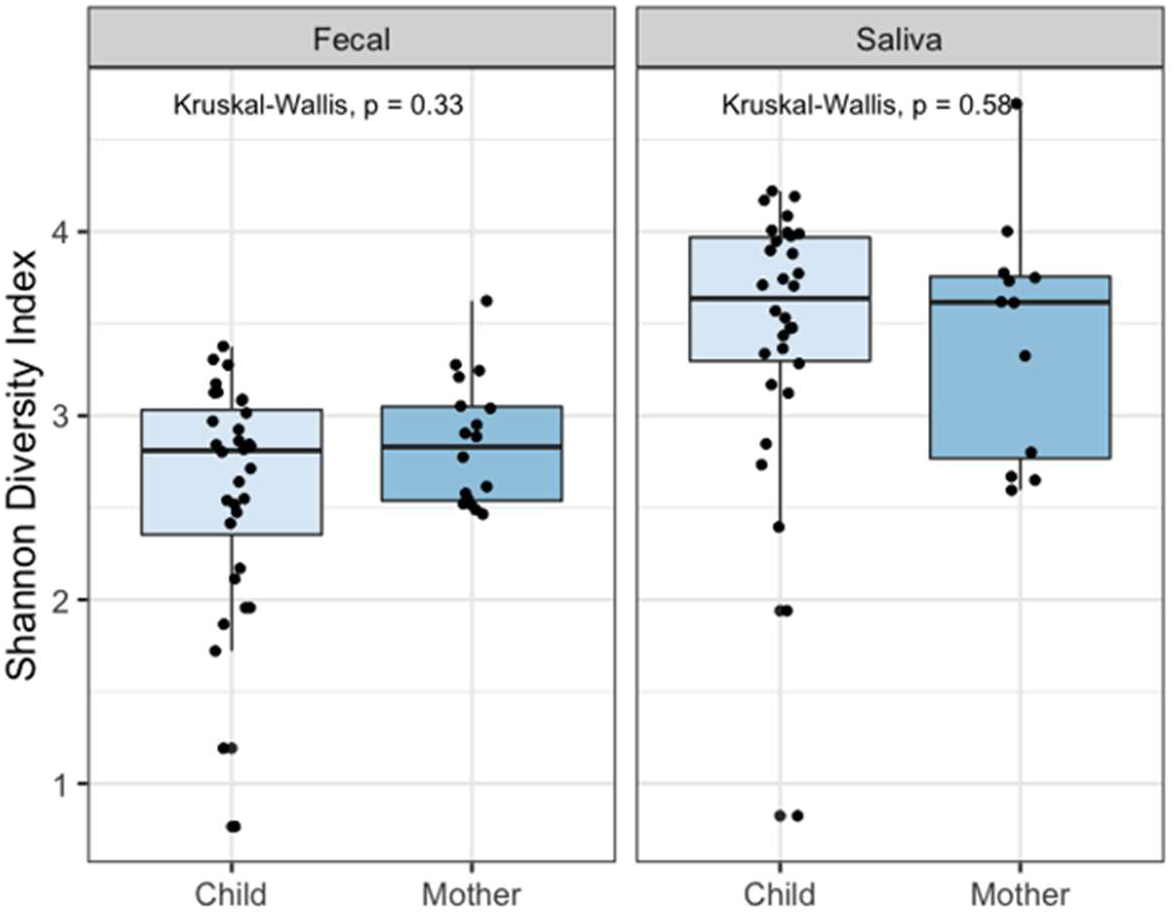
Comparison of microbial diversity between mother-offspring groups as measured by Shannon’s Diversity Index. Comparison of sample diversity by the Kruskal-Wallis test showed that the median diversity did not significantly differ between mothers and offspring in either fecal or saliva samples (p = 0.33 and p = 0.58 respectively).

In order to determine if there was a difference between the microbiome composition of household units in the village and the greater community at large the β-diversity in fecal and saliva samples was calculated using the Bray Curtis Dissimilarity measure both within households and between households (**Fig. 4**).

**Figure 4.**
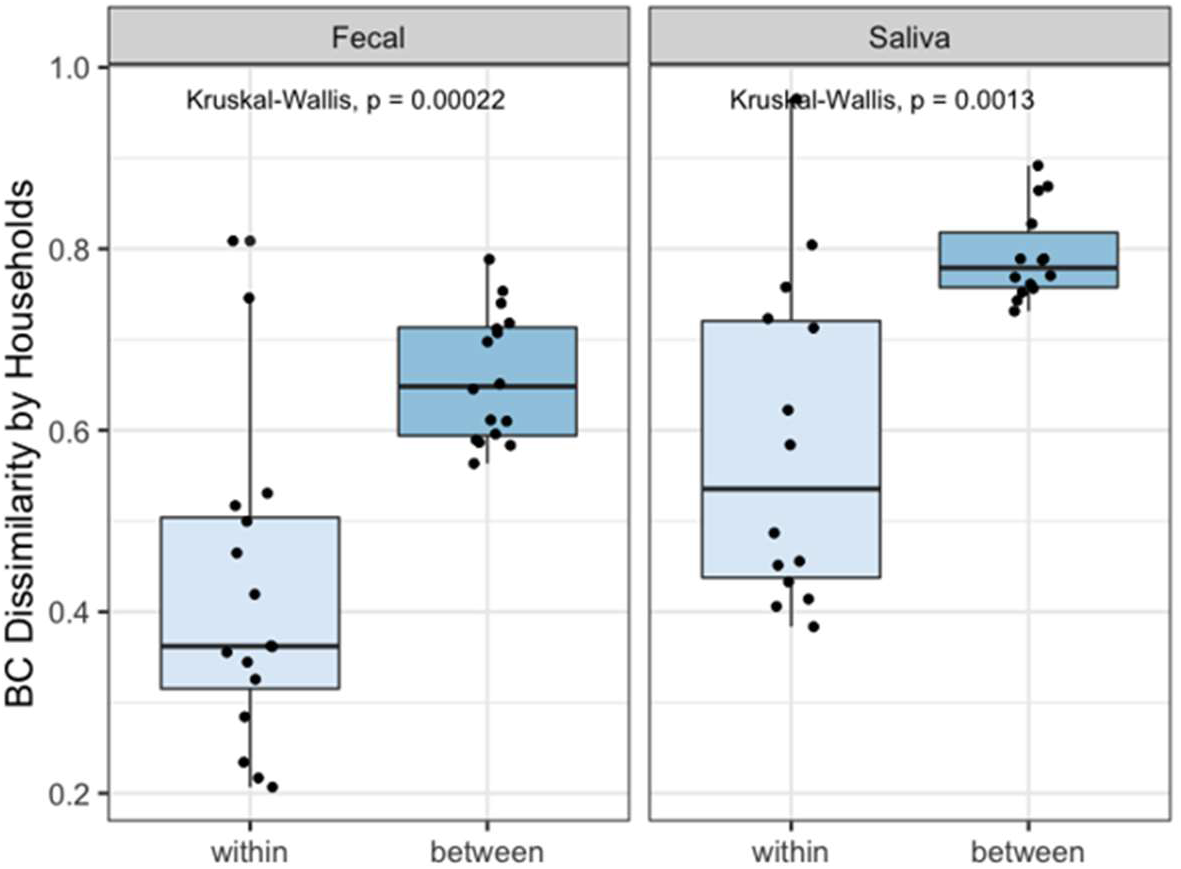
Comparison of dissimilarity within and between household units. Bray Curtis Dissimilarity measures were compared using the Kruskal-Wallis test, showing that members of a household were less dissimilar to each other than to the community at large. This difference in dissimilarity was significant in both fecal and salivary samples (p = 0.00022 and p = 0.0013 respectively).

Samples of feces and saliva from within the same household unit were found to have mean dissimilarities of 40.9% and 61.6%, respectively. Meanwhile, samples of feces and saliva from between different households in the community were found to have mean dissimilarities of 65.9% and 81.1%, respectively. The difference in dissimilarity between household units and the community was assessed using the Kruskal-Wallis test. The analysis determined that there is a significant difference between the dissimilarity of microbiomes within a household compared to the dissimilarity between households (p = 0.00022 for fecal samples, p = 0.0013 for saliva samples). The variation in microbiome composition of fecal and saliva samples was determined by PERMANOVA to be largely explained by the household from which they originate (R^2^ = 0.42, p = 0.001 and R^2^ = 0.67, p = 0.017 respectively). Residual variation was largely explained by the individual due to averaging of microbiome samples.

## DISCUSSION

In this study we showed, using two common and robust measures of microbial diversity, that the fecal and oral microbiomes of mother-offspring pairs in the indigenous Conambo river village were more similar to one another than to the microbiomes of their surrounding village community. This increase in the similarity of microbiomes between mother-offspring groups has been posited to be due to the formative influence of the mother on infant microbiome development.^2^ The results of this study indicate that a large percentage of the variation in diversity within the microbiomes of this isolated community can be explained by the household in which they live. This household variation can likely be attributed to the shared proximity, microenvironment, and specific daily diet of each family.^1,3–5^ While this initial investigation was not sufficient to conclude a causal relationship of the microbial transfer from mother to child it did present evidence to support further investigation and research into this area. Prior research has investigated the relative influence of host genetics and environmental conditions on the development and composition of the human microbiome, showing that environment was a large influence on the microbiome.^3^

This study presented the preliminary analysis on an isolated microbiome community in order to determine to what degree there are shared species or functional capacities of oral and fecal microbiomes between family members living together, namely maternal-offspring groups. Additional analysis will involve examining the roles of specific shared species or strains which are present in the different microbiomes as well as annotation of the functional genes present, which will help to indicate metabolic processes encoded by these microbes. Furthermore, analysis of shared functions and strains between mother-offspring oral microbiomes and the chicha microbiomes from that household. The motivation for such an investigation is the maternal nurturing practice within this community, where mothers wean their children off of breast milk at relatively young ages by feeding them the pre-masticated yuca pulp which is fermented into chicha.^11,12^ This weaning practice is not unique to this village and has been reported in another study which suggested it as a vector for microbial transfer between mothers and children.^2^

There is also a great deal of anthropological work to still take place alongside microbial analysis. Additional ethnographic and detailed individual data can be gathered and compiled to increase the accuracy of computational analysis. Attempts to gather culturable samples of microbes from the indigenous people could also lead to a great number of discoveries. A prominent issue with this study was that a large majority of the sequences in this dataset could not be identified by currently available databases. This issue of underrepresentation or absence of comparison sequences in databases is a continuous problem in microbiome studies, showing that there still needs to be a great deal of effort in culture and characterization of novel microbial species from unique or isolated cohorts. This issue has been observed by other researchers and there are now attempts to characterize and preserve distinct microbiomes from around the world.^16^ In future studies with this dataset the use of additional databases and sequence annotation pipelines would provide greater clarity and insight into this indigenous group.

## ACKNOWLEDGEMENTS

I would first and foremost like to thank Andrew Oliver for his all of time and effort in helping with the statistical analysis and figure generation in R. I would also like to thank Dr. Katrine Whiteson and all the wonderful researchers in the Whiteson lab for their help and guidance through this project. An incredible amount of praise is due to Dr. John Patton who works with the people of Conambo in Ecuador. Without Dr. Patton’s dedicated work over the decades any research on this indigenous group would not be possible. Finally, thanks to Dr. Patton’s grad students, especially my fiancée Alexandria, who all went out into the field with him, spending a month in the remote jungle to collect and catalogue the biological samples. I would also like to acknowledge the UCI Microbiome Initiative, who supported the sequencing for this project with a pilot award.

## DATA AND SEQUENCE AVAILABILITY

Raw sequencing data, analysis code, and other raw data or supplementary material will be made available on public databases, pending further peer reviewed publications.

